# Age-dependent changes in electrophysiology and calcium handling – implications for pediatric cardiac research

**DOI:** 10.1101/657551

**Authors:** Luther M. Swift, Morgan Burke, Devon Guerrelli, Manelle Ramadan, Marissa Reilly, Damon McCullough, Ashika Chaluvadi, Colm Mulvany, Rafael Jaimes, Nikki Gillum Posnack

## Abstract

**Rationale:** The heart continues to develop and mature after birth and into adolescence. Accordingly, cardiac maturation is likely to include a progressive refinement in both organ morphology and function during the postnatal period. Yet, age-dependent changes in cardiac electrophysiology and calcium handling have not yet been fully characterized.

**Objective:** The objective of this study, was to examine the relationship between cardiac maturation, electrophysiology, and calcium handling throughout postnatal development in a rat model.

**Methods:** Postnatal rat cardiac maturation was determined by measuring the expression of genes involved in cell-cell coupling, electrophysiology, and calcium handling. In vivo electrocardiograms were recorded from neonatal, juvenile, and adult animals. Simultaneous dual optical mapping of transmembrane voltage and calcium transients was performed on isolated, Langendorff-perfused rat hearts (postnatal day 0–3, 4-7, 8-14, adult).

**Results:** Younger, immature hearts displayed slowed electrical conduction, prolonged action potential duration and increased ventricular refractoriness. Slowed calcium handling in the immature heart increased the propensity for calcium transient alternans which corresponded to alterations in the expression of genes encoding calcium handling proteins. Developmental changes in cardiac electrophysiology were associated with the altered expression of genes encoding potassium channels and intercalated disc proteins.

**Conclusion:** Using an intact whole heart model, this study highlights chronological changes in cardiac electrophysiology and calcium handling throughout postnatal development. Results of this study can serve as a comprehensive baseline for future studies focused on pediatric cardiac research, safety assessment and/or preclinical testing using rodent models.

## Introduction

The mammalian heart begins as a mesoderm derived tube with slow electrical conduction, an underdeveloped sarcoplasmic reticulum and limited contractile force (van Weerd & Christoffels, 2016; Günthel *et al.*, 2018). The mammalian heart continues to develop postnatally, not reaching maturity in small rodents until weeks after birth (Vreeker *et al.*, 2014). Because of this extended period of maturation, the juvenile heart is uniquely dynamic and often considered a ‘moving target’ (Pesco-Koplowitz *et al.*, 2018). As such, there are considerable gaps in our understanding of normal cardiac physiology throughout the postnatal period. These gaps in knowledge can have unintended consequences, particularly when pharmacological therapies or toxicological studies are designed and tested using adult cardiac models – targeting ion channels and/or signaling pathways that may be underdeveloped in the immature myocardium. Indeed, a number of age-dependent cardiac responses to antiarrhythmics have been reported (e.g., dofetilide (Obreztchikova *et al.*, 2003), sotalol (Saul *et al.*, 2001)) and inotropes (e.g., dobutamine, isoproterenol (Driscoll *et al.*, 1980)). Likewise, the Cardiac Safety Research Consortium recently highlighted the need for research studies focused on developmental cardiac physiology, with the goal of improving cardiac safety testing. Specifically, the consortium emphasized a need for “functional, systematic and comprehensive studies of cardiac development” to bolster pediatric cardiac research (Bates *et al.*, 2012).

Small rodent animal models are frequently employed in cardiovascular research, as the rodent and human heart follow a similar sequence of structural cardiac development and electrophysiological patterning (Marcela *et al.*, 2012; Krishnan *et al.*, 2014; van Weerd & Christoffels, 2016). As with humans, the rodent heart exhibits electrical restitution properties and an action potential that is rate-dependent (Knollmann *et al.*, 2007). Immature cardiac myocytes from both humans and rodents lack fully developed transverse tubules and sarcoplasmic reticulum (Ziman *et al.*, 2010), which influences calcium cycling (Escobar *et al.*, 2004; Wagner *et al.*, 2005) and contractile function (Louch *et al.*, 2015; Racca *et al.*, 2016). Additionally, immature rodent and human cardiomyocytes have underdeveloped intercalated discs, which impacts cell-cell coupling and electrical conduction (Vreeker *et al.*, 2014). Moreover, age-dependent changes in voltage-gated potassium (Wahler *et al.*, 1994; Wang & Duff, 1997; Grandy *et al.*, 2007), calcium (Wang *et al.*, 2003; Wagner *et al.*, 2005) and sodium channel (Cai *et al.*, 2011) current have been described. Although species-specific characteristics do exist; for instance an age-dependent increase in the heart rate of small rodents (Heier *et al.*, 2010) is coupled to changes in excitation-contraction coupling that allow the rodent heart to adapt to faster beating rates. Accordingly, the rodent action potential (measured via intracellular microelectrode) becomes progressively shorter as cardiomyocytes mature (Escande *et al.*, 1985; Wahler *et al.*, 1994; Wang *et al.*, 2003). Nevertheless, small rodents remain a vital tool to investigate cardiac maturation, electrophysiology and excitation-contraction coupling (Krishnan *et al.*, 2014; Günthel *et al.*, 2018).

The objective of this study was to evaluate the relationship between cardiac maturation, electrophysiology and calcium handling throughout postnatal rat development. To date, developmental changes in rodent cardiac electrophysiology and calcium handling have largely been limited to cell models (Artman *et al.*, 2000). However, the results gleaned from these models may not translate to a whole heart with specialized anatomy, cell populations (atrial, nodal, ventricular), spatial tissue heterogeneity, and a coordinated conduction system. Therefore, we utilized a three-dimensional whole heart model to describe temporal changes in rat cardiac electrophysiology and calcium handling during postnatal development. Simultaneous, dual-optical mapping of transmembrane voltage (V_m_) and calcium transients (Ca^2+^) was performed on isolated Langendorff-perfused hearts. The results of this study can serve as a baseline for future pediatric cardiac research studies, focused on environmental, pharmacological or toxicological perturbations.

## Methods

Animal protocols were approved by the Institutional Animal Care and Use Committee at Children’s Research Institute and followed the Nationals Institutes of Health’s *Guide for the Care and Use of Laboratory Animals*. Experiments were performed using Sprague-Dawley rats from postnatal day (PND) 0 to adulthood (2-3 months), Taconic Biosciences (n = 126). Animals were housed in conventional acrylic rat cages in the Research Animal Facility, under standard environmental conditions (12:12 hour light:dark cycle, 64 – 78C, 30-70% humidity, free access to reverse osmosis water, corn cob bedding and #2918 rodent chow, Envigo).

### In vivo surface electrocardiogram recordings

Electrocardiograms (ECG) were collected from conscious animals using an ecgTUNNEL system (emka Technologies). The platform electrodes were coated with ultrasound gel prior to placing the animal in the system. A clear half-tunnel was carefully positioned over the top of older animals (>pnd 6), to limit movement which can introduce noise in the signal. The animals were acclimated to the platform for 5 min, biosignals were continuously acquired for 2 min using iox2 (emka Technologies), and ECG segments were computed in ecgAUTO (emka Technologies).

### Isolated heart preparation and electrophysiology measurements

Animals were anesthetized with 2% isoflurane; the heart was rapidly excised, and the aorta cannulated. The heart was then transferred to a temperature-controlled (37°C) constant-pressure (70 mmHg) Langendorff perfusion system. Excised hearts were perfused with Krebs-Henseleit buffer bubbled with 5% CO^2^ and 95% oxygen throughout the duration of the experiment, as previously described (∼1 hour) (Jaimes III *et al.*, 2014). Electrocardiograms were recorded throughout the duration of the experiment in a lead II configuration. During sinus rhythm, ECG signals were collected to analyze heart rate, atrial depolarization, atrioventricular conduction (PR interval), ventricular depolarization time (QRS width), and heart rate variability, including root means successive square difference (rMSSD) and standard deviation of the normal RR intervals (SDNN). Signals were acquired in iox2 and ECG parameters were quantified in ecgAUTO.

### Optical mapping

To reduce motion artifact during imaging experiments, the heart was perfused with Krebs-Henseleit buffer supplemented with 10 μM (-/-) blebbistatin (Sigma-Aldrich) (Fedorov *et al.*, 2007; Swift *et al.*, 2012). Epicardial imaging was performed by sequentially loading the heart with fluorescent dyes through a bubble trap located proximal to the aortic cannula (Kay *et al.*, 2008; Posnack *et al.*, 2014*a*). A calcium indicator dye (50 μg Rhod2-AM) (Lang *et al.*, 2011; Jaimes *et al.*, 2016*a*) was added and allowed to stabilize for 10 min, followed by a potentiometric dye (62.1 μg RH237) (Swift *et al.*, 2008). The epicardium was illuminated with an LED spotlight (530 nm, 200 mW; Mightex), fitted with an optical filter (ET530/40x nm, Chroma Technologies). Fluorescence signals were acquired using an image splitting device (Optosplit II, Cairn Research Ltd) positioned in front of a sCMOS camera (Zyla 4.2Plus, Andor Technologies). The path splitter was configured with a dichroic mirror (660+nm, Chroma Technologies) that passed RH237 emission and reflected Rhod-2AM fluorescence. RH237 fluorescence was longpass filtered (ET710, Chroma Technologies) and Rhod-2AM was bandpass filtered (ET585/40m, Chroma Technologies). A fixed focal length 17mm/F0.95 lens was attached to the image splitting device (Schneider, #21-010456). *MetaMorph* (Molecular Devices LLC) was used for optosplit image alignment and LED on/off triggering. Transmembrane voltage and calcium signals were acquired simultaneously at 800 frames per second.

For ventricular pacing, a 0.25 mm diameter tungsten, unipolar, cathodal electrode was placed on the left ventricle’s epicardium, and a stainless-steel indifferent electrode was placed under the heart. To determine the ventricular effective refractory period (VERP), dynamic pacing (S1-S1) was performed during optical mapping. A Bloom Classic electrophysiology stimulator (Fisher Medical) was set at a pacing current 1.5x the minimum pacing threshold (∼1.8 mA) with 1 msec monophasic pulses; pacing cycle length (PCL) was decremented stepwise (250 – 50 msec) until a loss of capture was observed, in order to identify VERP.

### Signal Processing

Following image acquisition, signal processing and data analysis were performed using a custom MATLAB script, as previously described (Posnack *et al.*, 2014*b*; Jaimes *et al.*, 2016*b*). A region of interest (appx 0.75 mm) was selected in identical locations on the split image of the heart for each raw image. The signals from each ROI corresponding to transmembrane voltage and calcium transients, were independently averaged and plotted against time. A peak detector algorithm was applied, and characteristics of each waveform were measured and averaged, including: action potential duration at 30% (APD30) and 80% (APD80) repolarization, APD triangulation (APD80-APD30) (Hondeghem *et al.*, 2001), and calcium transient duration at 30% (CaD30) and 80% (CaD80) reuptake. Optical signals were acquired during each pacing cycle length (PCL). Calcium transient alternans were defined as sequential calcium transient measurements that differed by >5%.

### Gene Expression Analysis

Total RNA was isolated from heart tissue using a RNeasy fibrous tissue kit (Qiagen). RNA was reverse transcribed using a SuperScript VILO cDNA kit (Thermo Scientific), and Taqman gene expression assays (Thermo Scientific) were used for quantitative real-time PCR (qPCR) analysis via a Quantstudio 7 platform (Applied Biosystems). Relative gene expression was assessed by normalizing CT values (dCT) to the housekeeping gene glyceraldehyde-3-phosphate dehydrogenase (*GAPDH)*, wherein a dCT >0 indicates reduced expression relative to the housekeeping gene and dCT <0 indicates increased expression relative to the housekeeping gene. dCT values allow for comparison between genes of interest, as compared to fold-change measurements. dCT values are reported as an average of technical replicates, with each assay including a minimum of 3 individual biological replicates.

Genes of interest include: myosin heavy chain 7 (*MYH7*), myosin heavy chain 6 (*MYH6*), gap junction protein α1 (*GJA1*), desmoplakin (*DSP*), junctophilin 2 (*JPH2*), junction plakoglobin (*JUP*), n-cadherin (*CDH2*), caveolin-3 (*CAV3*), tight junction protein 1 (*TJP1*), SERCA Ca^2+^-ATPase (*SERCA2*), calsequestrin 2 (*CASQ2*), ryanodine receptor 2 (*RYR2*), sodium-calcium exchanger 1 (*SLC8A1*), voltage-dependent T-type calcium channel α-1G (*CACNA1G*), voltage-gated potassium channel subunit Kv4.2 (*KCND2*), voltage-gated potassium channel subunit Kv1.5 (*KCNA5*), voltage-gated potassium channel subunit Kv4.3 (*KCND3*), and cardiac inward rectifier potassium channel (*KCNJ2*).

### Statistical Analysis

Data are presented as mean ± standard deviation. Statistical analysis was performed using one or two-way analysis of variance and false discovery rate (0.1) to correct for multiple comparisons testing using GraphPad Prism. Significance was defined as (*p≤0.05). Optical signals were analyzed using custom algorithms (MATLAB). Optical signals were collected for 2 seconds per pacing frequency, resulting in 8 – 25 signals per group.

## Results

### Postnatal heart development and in vivo electrophysiology

The postnatal heart maintains limited proliferative potential, progressing from hyperplasia to hypertrophic growth shortly after birth (**Figure 1A-D**) (Li *et al.*, 1996). In rodents, this time course is associated with a shift in myosin heavy chain expression (*MYH7* to *MYH6*) and an age-dependent increase in the spontaneous beating rate (Mahdavi *et al.*, 1984). We observed a similar shift in expression, compared to immature hearts collected on postnatal day 0-3 (PND 0-3), MYH7 expression decreased by 96% and 155% in older hearts (PND 4-10, PND >10, respectively). Whereas MYH6 expression increased by 56% and 222% in older hearts (PND 4-10, PND >10, respectively) compared with PND 0-3, **Figure 1D**. Myosin heavy chain isoform expression complements changes in heart rate, as MYH6 kinetics are three times faster than MYH7 (Galler *et al.*, 2002). As cardiac development progressed, the temporal change in MYH expression corresponded with a linear increase in the *in vivo* resting heart rate from 181 BPM (PND 0) to 429 BPM (PND 10), **Figure 2B**. No significant difference in resting heart rate was observed in animals older than PND 10. Continued development of autonomic influences (Robinson, 1996) on the heart were measured via time-domain indices of heart rate variability. Compared to the earliest time point (PND 0), standard deviation of heart rate (SDNN) increased by 347% and 849% in older animals (PND 10 and adult, respectively), **Figure 2C**. Similarly, the root means successive square difference (rMSSD) increased by 145% and 689% in older animals (PND 10 and adult, respectively) compared with the earliest time point measured (PND 0), **Figure 2D**.

**Figure 1.**
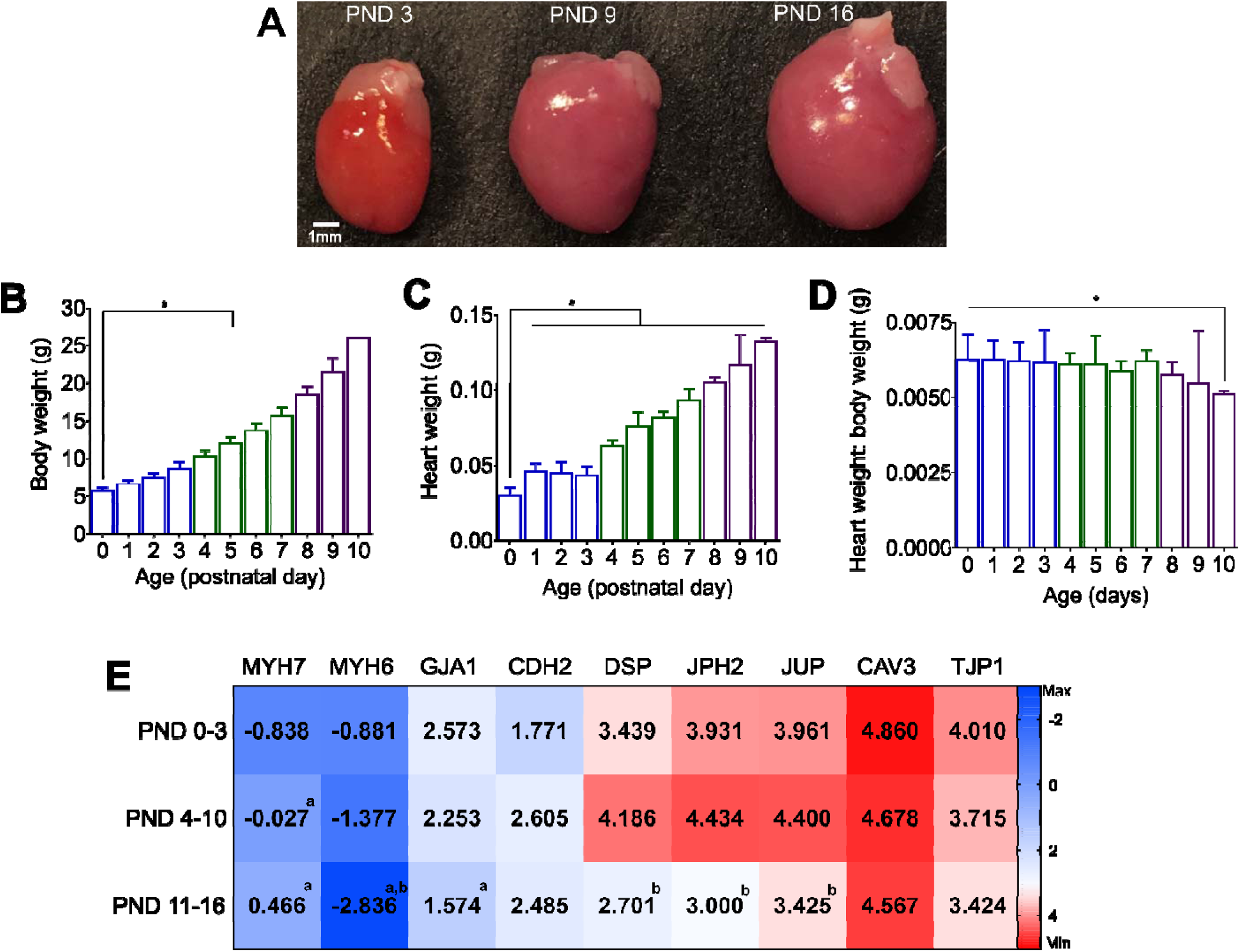
Postnatal development and cardiac maturation. **(A**) Isolated whole hearts from postnatal day 3, 9 and 16. (**B)** Age-dependent increase in body weight and (**C)** heart weight, and (**D)** slight decrease in the heart weight to body weight ratio. (**E)** Developmental time course corresponds to shift in myosin heavy chain gene expression (*MYH7* to *MYH6*), and increased expression of key genes involved in intercellular coupling via gap junctions (*GJA1*) and desmosomes (*DSP, JPH2, JUP*). Gene expression scale depicts maximal expression across all genes in this cohort (blue) and minimal expression (red), relative to *GAPDH* housekeeping gene. PND = postnatal day; ‘a’ denotes significant difference relative to PND 0-3 age, ‘b’ denotes significant difference relative to PND 4-10 age. mean ± SD, n≥3 independent experiments.

**Figure 2.**
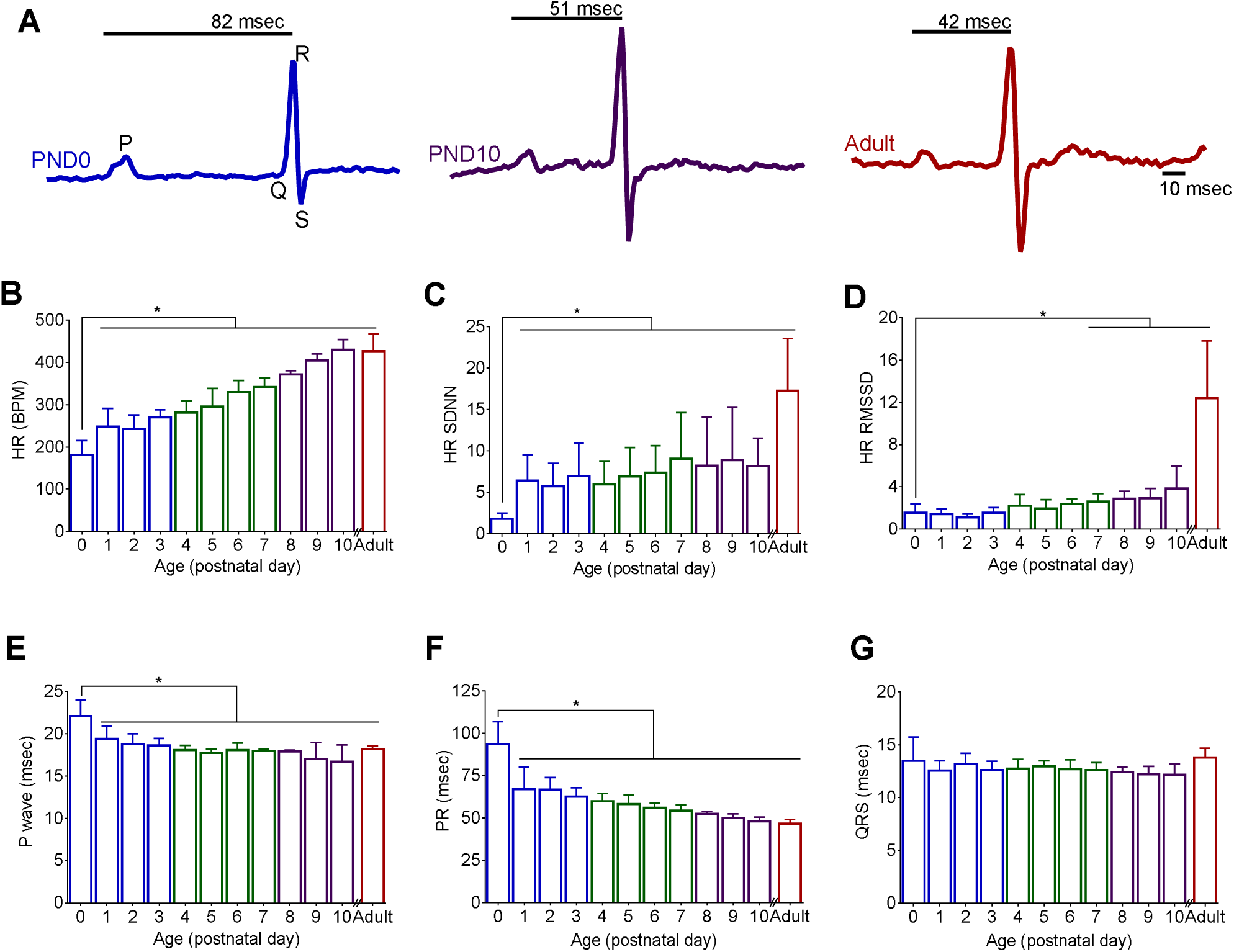
Age-dependent alterations in *in vivo* electrocardiogram parameters. **(A)** Example of non-invasive electrocardiogram waveforms recorded from postnatal day 0 (PND 0), 10 (PND 10) and adult animal; PR interval time is denoted. Postnatal development was associated with an age-dependent increase in heart rate (**B)**, heart rate variability (**C,D)**, shortening of p-wave duration (**E)** and shortening of the PR interval (**F)**. No significant difference in the QRS interval time was observed using non-invasive recordings (**G)**. *indicates statistically significant difference from earliest measured time point (PND 0). PND = postnatal day, n≥7 animals per age, mean ± SD, *p<0.05

Along with an age-dependent increase in heart rate, we observed a progressive shortening of *in vivo* electrocardiogram parameters during sinus rhythm. Atrial conduction (P-wave duration) decreased from 22.1 msec (PND 0) to 16.6 msec (PND 10), and atrioventricular conduction time (PR interval) shortened from 93.8 msec (PND 0) to 48.1 msec (PND 10), **Figure 2A,E,F**. An age-dependent trend in ventricular depolarization time (QRS interval) using non-invasive electrocardiogram monitoring was not observed (**Figure 2G)**. Age-dependent shortening of atrial and atrioventricular conduction can be partly attributed to remodeling of cardiomyocyte size (Spach *et al.*, 2000) and more defined intercellular connections (Vreeker *et al.*, 2014), which have both been shown to enhance electrical propagation. Compared with immature hearts (PND 0-3), older hearts (PND >10) had an increased expression of cell coupling genes encoding gap junction (39% GJA1) and desmosomal proteins (99% DSP, 24% JPH2, 14% JUP), **Figure 1D**.

### Age-dependent shortening of action potential duration and refractory period

Compared with neonatal cardiomyocytes, adult cells have increased potassium channel expression that can expedite repolarization and shorten action potential duration (APD) time (Escande *et al.*, 1985; Wahler *et al.*, 1994; Wang *et al.*, 2003). To evaluate action potential shape and duration in the whole heart, transmembrane voltage signals were recorded from the epicardial surface of isolated, Langendorff-perfused hearts. Optical signals acquired from juvenile hearts were binned into three age groups (PND 0-3, PND 4-7, and PND 8-14) and compared with adults. Prolonged APDs were consistently observed in younger hearts at multiple pacing frequencies. At 250 msec PCL, APD30 shortened from 54.4 msec to 20 msec and APD80 shortened from 113.2 msec to 60.5 msec (PND 0-3 vs adult, respectively), **Figure 3A-C.** At faster pacing rates (150msec PCL), APD30 and APD80 prolongation was observed in younger hearts, albeit the difference between intermediate age groups was modest. Additionally, triangulation of action potential shape was more pronounced in younger hearts (58.4 msec, PND 0-3) compared to older hearts (30.4 msec, adult), **Figure 3D.**

**Figure 3.**
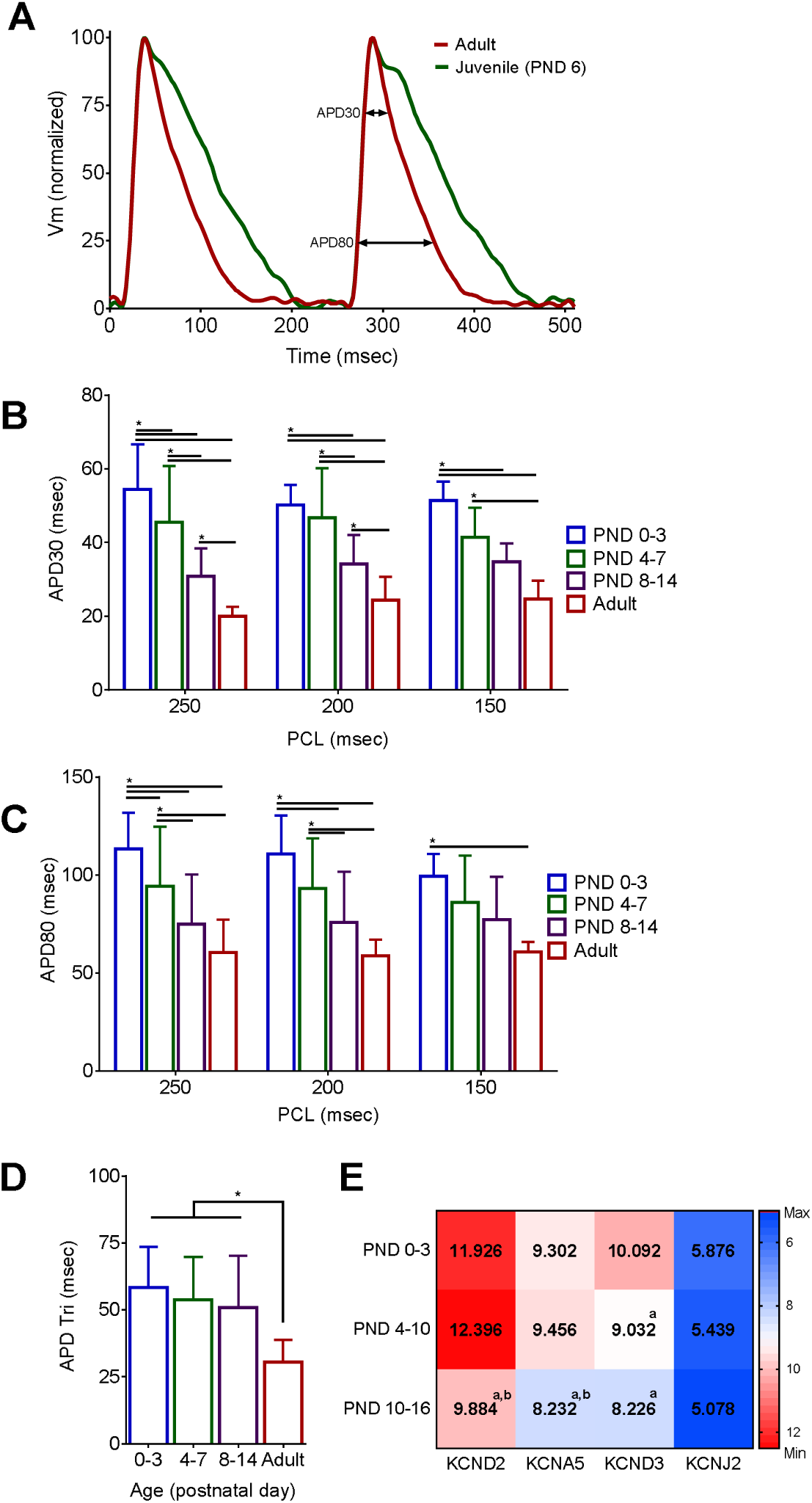
Age-dependent shortening of action potential duration time. **(A)** Transmembrane voltage (Vm) signals optically mapped from the epicardial surface of excised, intact hearts (250 msec PCL). (**B,C)** Prolonged APDs were observed in younger hearts (APD30, APD80), which also displayed more triangulated action potentials (**D)**. Shortened APD in older animals coincided with an age-dependent increase in voltage-gated potassium channel gene expression (**E)**. Data binned into the following age groups: PND 0-3, PND 4-7, PND 8-14 and adult (2-3months). n≥5 individual hearts per age. APD30 = action potential duration at 30% repolarization, APD80 = 80% repolarization, APD Tri = Triangulation APD80-APD30, PND = postnatal day. n≥7 animals per age, mean ± SD, *p<0.05

Increased potassium channel current can be mechanistically linked to shorter, less triangulated action potentials (Grandy *et al.*, 2007). We observed an age-dependent increase in potassium channel gene expression during postnatal cardiac maturation. This included a 17% increase in KCND2 and 18% increase in KCND3 gene expression (PND 0-3 vs adult hearts), which corresponds to the fast transient outward It**_0_** current (**Figure 3F)**. Importantly, It**_0_** is responsible for the rapid repolarization and lack of a plateau phase in the adult rodent action potential (Wang & Duff, 1997; Knollmann *et al.*, 2007). To further investigate the relationship between potassium channel expression and action potential duration time in the immature heart, an epicardial pacing protocol was implemented to pinpoint the ventricular effective refractory period (VERP). The VERP shortened with increasing age, from 183 msec in younger hearts (PND 0-3) to 115 msec in adult hearts (**Figure 4A-C**). With increased ventricular refractoriness in younger hearts, loss of capture at PCLs <150 msec prevented APD measurements at faster pacing frequencies.

**Figure 4.**
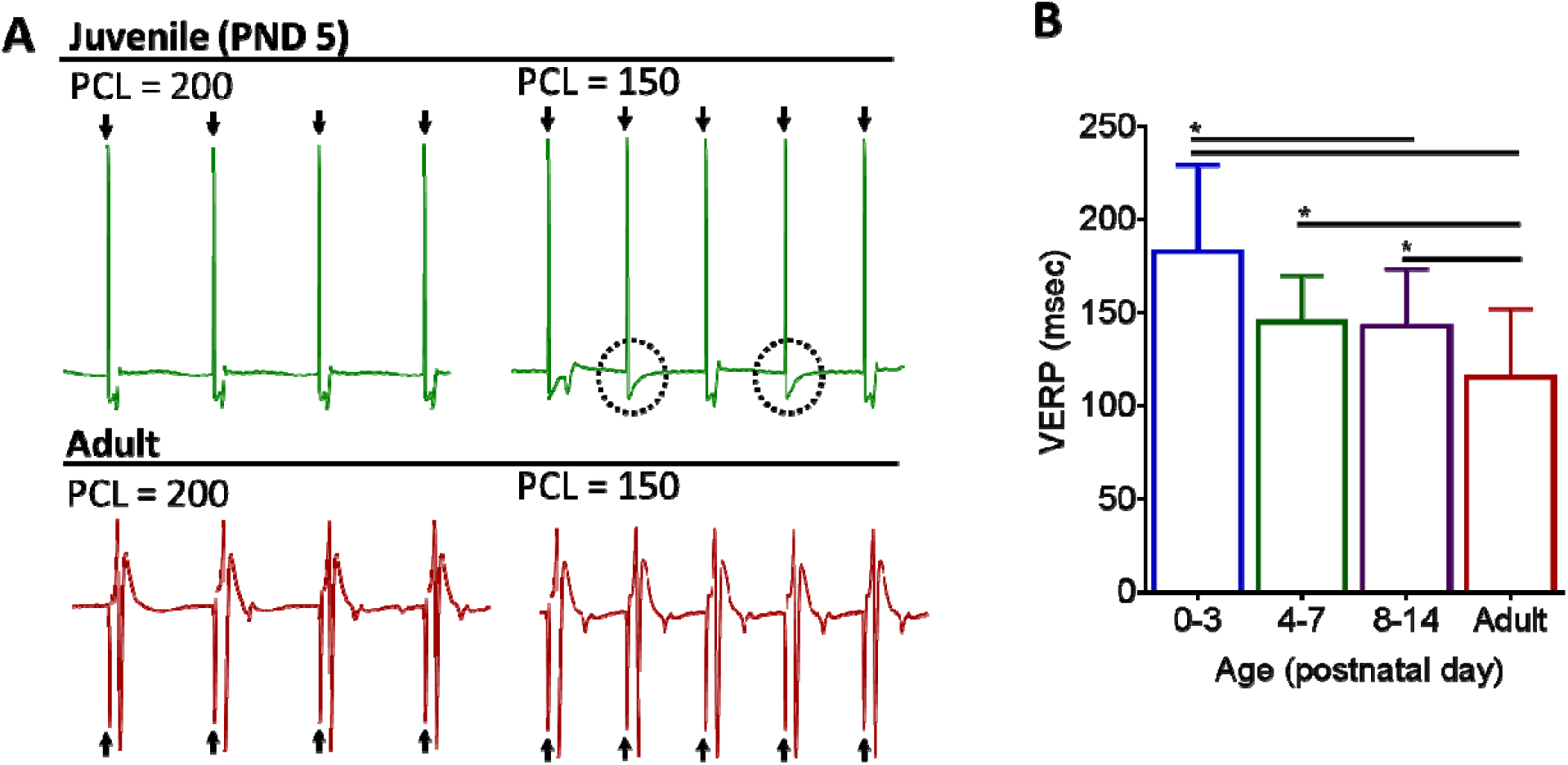
Age-dependent shortening of the ventricular effective refractory period (VERP) **(A)** Excised hearts were paced near the apex at two pacing cycle lengths (PCL = 250 msec, 150 msec) and electrocardiograms were recorded. Pacing spikes denoted with arrows. Top: Juvenile heart at both PCL. Note the loss of capture, circled, at a shorter cycle length. Bottom: Adult heart shows ventricular response to each pacing spike, at both pacing frequencies. (**B)** Age-dependent decrease in the VERP. n≥5 animals per age, mean ± SD, *p<0.05

### Age-dependent alterations in calcium handling and incidence of alternans

Cardiomyocyte maturation includes the invagination of transverse tubules, formation of couplons and synchronized calcium-induced calcium release (Ziman *et al.*, 2010). To evaluate calcium handling in the whole heart, calcium transients were recorded from the epicardial surface of isolated Langendorff-perfused hearts. Rate adaptation of the calcium transient duration time was observed in all age groups. For example, in the PND0-3 age group CaD80 decreased from 168 to 151 to 133 msec (PCLs = 250, 200, 150msec). But, immature hearts had consistently slower calcium handling at each pacing cycle. At slower pacing frequencies (250 msec PCL), CaD30 shortened from 95 msec to 57 msec and CaD80 shortened from 168 msec to 115 msec (PND 0-3 vs adult), **Figure 5A-C**. At faster pacing cycles (150 msec), CaD30 and CaD80 were longer in younger hearts, but differences between age groups were modest.

**Figure 5.**
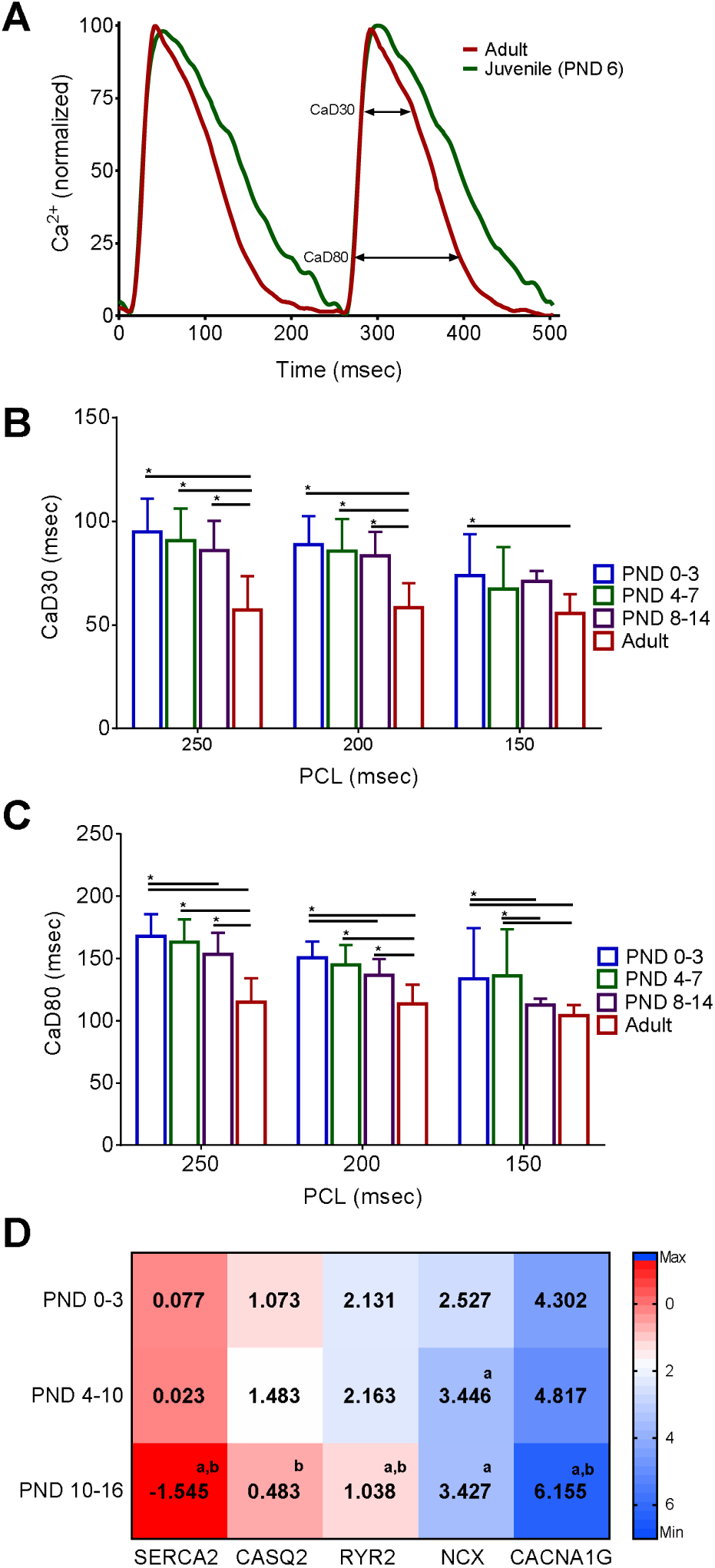
Age-dependent shortening of calcium handling. **(A)** Calcium transients recorded from the epicardial surface of excised, intact hearts (250 msec PCL). (**B,C)** Prolonged CaDs were observed in younger hearts (CaD30, CaD80). Faster calcium handling in older animals coincided with an age-dependent increase in key calcium handling genes associated with calcium binding in the sarcoplasmic reticulum (CASQ2), calcium release (RYR), calcium reuptake into the SR (SERCA2) and an age-dependent decrease in the sodium-calcium exchanger (NCX) and immature T-type calcium channels (CACNA1G). Data binned into the following age groups: PND 0-3, PND 4-7, PND 8-14 and adult (2-3 months). n≥5 individual hearts per age. CaD30 = calcium transient duration at 30%, CaD80 = 80%, PND = postnatal day. n≥7 animals per age, mean ± SD, *p<0.05

Faster calcium handling in adult hearts was linked to an age-dependent increase in genes associated with calcium binding within the sarcoplasmic reticulum (55% increase CASQ2 vs PND 0-3), calcium release (49% increase RYR2 vs PND 0-3), and calcium-reuptake into the sarcoplasmic reticulum (SERCA2), **Figure 5D**. Whereas immature cardiomyocytes rely less on calcium-induced calcium release, and more on sarcolemma calcium influx via the sodium-calcium exchanger (NCX) and T-type calcium channels (CACNA1G) (Louch *et al.*, 2015). In older hearts, we observed a 36% decrease in NCX expression and 43% decrease in CACNA1G compared with hearts aged PND 0-3, **Figure 5D**. Disturbances in calcium handling have also been associated with an increased incidence of alternans, or beat-to-beat alterations in calcium transient amplitude and kinetics (Edwards & Blatter, 2014; Ramadan *et al.*, 2018). A dynamic pacing protocol was implemented to pinpoint the alternans threshold, or, the longest pacing cycle length required to elicit calcium transient alternans. Immature hearts displayed an increased propensity for calcium transient alternans at slower pacing frequencies (160 msec, PND 0-3) compared with adult hearts (99 msec), **Figure 6A, B.**

**Figure 6.**
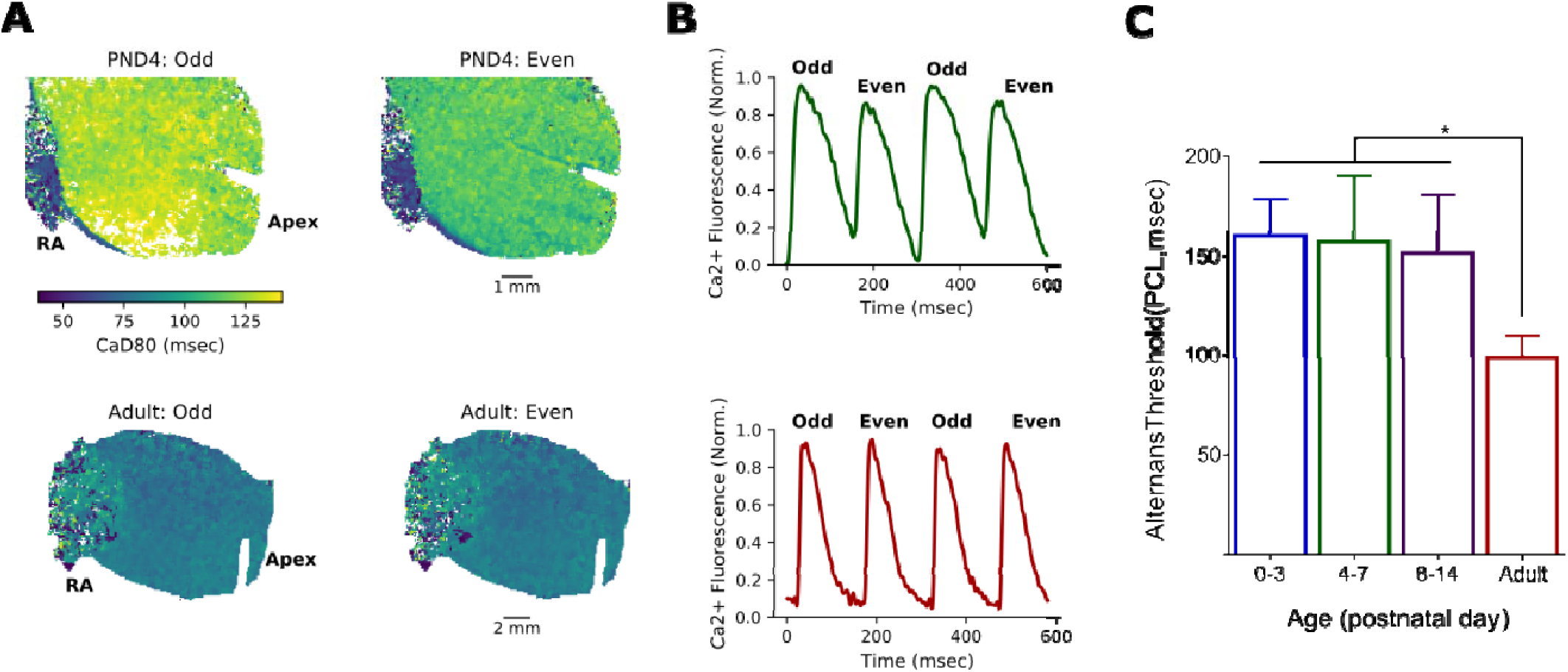
Increased incidence of calcium transient alternans in immature hearts. **(A)** Immature hearts (top two panels) displayed an increased susceptibility to calcium transient alternans compared with adult hearts (bottom two panels). Images show peak fluorescence at 150 msec PCL. (**B)** Green and red traces represent four paced beats from the neonatal (PND4) and adult heart, respectively. (**C)** The slowest PCL that resulted in alternating calcium transients (alternans threshold) was significantly slower in younger hearts compared with adults. Data binned into the following age groups: PND 0-3, PND 4-7, PND 8-14 and adult (2-3months). PND = postnatal day. n≥6 animals per age, mean ± SD, *p<0.05

## Discussion

Cardiac excitation-contraction coupling is the process by which an action potential evokes an increase in intracellular calcium, which subsequently triggers contraction (for review(Bers, 2001)). As the heart continues to develop postpartum, cardiac electrophysiology and excitation-contraction coupling dynamics mature with age. This occurs around 21 days postnatal in rats (van Weerd & Christoffels, 2016) and 20 years postnatal in humans (Mollova *et al.*, 2013). In this study, we examined the relationship between postnatal age, cardiac electrophysiology and calcium handling using *in vivo* and *in situ* models. We show that the immature heart displays slowed atrioventricular conduction, prolonged action potential duration time, and longer ventricular effective refractory periods, compared with adult hearts. Age-dependent alterations in cardiac electrophysiology were associated with changes in genes encoding voltage-gated potassium channels and intercalated disc proteins that facilitate intercellular coupling and electrical propagation. Calcium handling was slowed in the immature heart, which coincided with less developed excitation-contraction coupling machinery and an increased propensity for calcium transient alternans.

### Postnatal changes in cardiomyocyte morphology

Postnatal cardiac maturation includes the formation of intercellular connections between neighboring cardiomyocytes, or intercalated discs. The cardiac intercalated disc includes the colocalization of adherens junctions, desmosomes, gap junctions, sodium and potassium channels – which facilitates the rapid transmission of electrical activity, initiating contractile forces between neighboring myocytes (Noorman *et al.*, 2009; Wang *et al.*, 2012; Scuderi & Butcher, 2017). Indeed, the development of intercellular low resistance pathways is vital to the heart’s ability to function as a highly coordinated syncytium. The spatial distribution of desmosomal, fascia adherens, and gap junction proteins shifts throughout postnatal development, from sporadically distributed to densely concentrated at the terminal ends of neighboring adult cardiomyocytes. In rodents, the intercalated discs are formed within the first 20 days after birth, but continue to develop well past maturity (Angst *et al.*, 1997). Conversely in humans, the process is gradual with the colocalizing ion channels, adherens junctions, and gap junctions not being apparent until 6-7 years after birth (Peters *et al.*, 1994; Vreeker *et al.*, 2014).

In the presented study, we reported an increase in the relative mRNA abundance of key genes involved in intercellular connections and communication via the intercalated discs. From postnatal day 0 to 16, older hearts had increased mRNA expression of connexin-43 (GJA1) typically localized to gap junctions, as well desmosomal genes, including desmoplakin (DSP), plakoglobin (JPH2) and junctophilin (JUP). Notably, well-established intercalated discs have a direct influence on electrical conduction, and gap junction modifiers shorten ECG parameters and decrease the propensity for cardiac alternans (Hsieh *et al.*, 2015). Similarly, we observed a progressive shortening of ECG parameters (p-wave, PR interval) with postnatal age that coincided with increased expression of intercalated disc genes.

### Postnatal changes in excitation-contraction coupling

The immature heart transitions from hyperplasia to hypertrophic growth shortly after birth (Li *et al.*, 1996; Louch *et al.*, 2015) and as the cardiomyocytes increase in size, transverse tubules begin to form and invaginate into the cell interior and the sarcoplasmic reticulum becomes more developed (Tanaka *et al.*, 1998). These morphological changes facilitate the formation of dyads and couplons, wherein ryanodine receptors and L-type calcium channels are in close proximity (Scriven *et al.*, 2013). Concomitant with these organizational changes, cardiomyocytes become less dependent on the sarcolemma calcium influx and more reliant on calcium-induced calcium release (Escobar *et al.*, 2004; Ziman *et al.*, 2010; Hamaguchi *et al.*, 2013). Ziman, et al. correlated the timing of t-tubule and couplon formation with improved excitation-contraction coupling in isolated cardiomyocytes aged 10 – 20 days (Ziman *et al.*, 2010). The authors showed that myocytes up to PND 10 lacked a t-tubule system, whereas the t-tubule system of cells isolated from PND 20 hearts were indistinguishable from adult myocytes.

In the presented study, we observed a similar developmental time course, with increased mRNA expression of calsequestrin, ryanodine and the sarcoplasmic reticulum calcium ATPase in hearts aged 10-16 days compared with 0-3 days. We also observed an age-dependent decrease in genes associated with sarcolemma calcium influx – namely the sodium-calcium exchanger and T-type calcium channel. Consequently, younger, immature hearts displayed prolonged calcium transient duration times and an increased propensity for calcium transient alternans. Importantly, calcium alternans can be associated with T-wave alternans and electrical instabilities (Clusin, 2003, 2008; Edwards & Blatter, 2014). Our results are in agreement with the work by Escobar, et al. which showed that ryanodine had a negligible effect on 2-day old neonatal cardiomyocytes compared with cells isolated from 3-week old juvenile animals (Escobar *et al.*, 2004). The latter indicates minimal involvement of calcium-induced calcium release in the excitation-contraction coupling of immature hearts.

### Postnatal changes in cardiac electrophysiology

Postnatal cardiac maturation in humans and rodents both include an increase in cell size, formation of intercalated discs, invagination of t-tubules and increased dependence on calcium-induced calcium release for excitation-contraction coupling. One of the inherent dissimilarities between species is an age-dependent increase in the heart rate of small rodents (Heier *et al.*, 2010), which necessitates a progressively shorter action potential as rodent cardiomyocytes mature (Escande *et al.*, 1985; Wahler *et al.*, 1994; Wang *et al.*, 2003). In the presented study, we describe a linear increase in the heart rate and shortening of ECG parameters in older vs younger animals. This time course corresponds with a shift in myosin heavy chain expression to MYH6, which has kinetics that are three times faster than MYH7 (Galler *et al.*, 2002). Although humans and rodents exhibit electrical restitution properties and an action potential that is rate-dependent, the rodent action potential lacks a plateau phase due to differences in outward potassium current (Knollmann *et al.*, 2007; Grandy *et al.*, 2007). We observed an age-dependent increase in the expression of voltage-gated potassium channels – namely KCND2 and KCND3 that encode Kv4.2 and Kv4.3 and facilitate It**_0_** current. Indeed, It**_0_** is responsible for >50% of total outward potassium current and the very short APD that is characteristic of the adult mouse myocardium (Wang *et al.*, 1996). In our study, action potentials showed rate dependency in all age groups, but immature hearts displayed longer action potential duration times and ventricular effective refractory periods.

### Limitations

The scope of our study was limited to the age-dependent effects on cardiac electrophysiology and calcium handling using a rat model, therefore differences with human physiology should be considered. Despite many similarities, species-specific differences in cardiac physiology exist between rodents and humans. Nevertheless, rodent models remain a valuable tool for understanding cardiac maturation, as an ‘ideal’ human cardiac research model does not currently exist. Differentiated human embryonic stem cells and induced pluripotent stem cells hold promise, but methodologies to reproducibly mature these derived myocytes are still a work in progress (Feric & Radisic, 2016; Ruan *et al.*, 2016; Tiburcy *et al.*, 2017; Ronaldson-Bouchard *et al.*, 2018) and cell-based models cannot fully replicate a three-dimensional whole heart. Further, since the presented study largely utilized an isolated heart model, optical mapping data reflect developmental changes in myocardial physiology without neuronal input.

## Conclusion

The mammalian heart continues to mature postnatally, with substantial developmental changes in cardiac electrophysiology and calcium handling within the first few weeks after birth. This study utilized in vivo recordings and an ex vivo heart model to characterize the developmental time course for electrophysiology and calcium handling from postnatal day 0 – 14 and compared with adults. Results of this study can serve as a baseline for future studies aimed at assessing environmental, pharmacological or toxicological perturbations.

## DISCLOSURES

Nothing to disclose.

## FUNDING

This work was supported by the National Institutes of Health (R00ES023477 & R01HL139472 to NGP), Children’s Research Institute and Children’s National Heart Institute.

## AUTHOR CONTRIBUTIONS

LS, MB, DG, MR^1^, MR^2^, DM, AC, CM, RJ and NGP performed experiments; LS, MB, DG, MR^1^, MR^2^, DM, AC, CM, RJ and NGP analyzed data; LS, MB, DG, MR^1^, RJ and NGP prepared figures; NGP drafted manuscript; LS, MB, RJ and NGP conceived and designed experiments; LS, MG, DG, MR^1^, MR^2^, DM, AC, CM, RJ and NGP approved manuscript. ^1^Manelle Ramadan, ^2^Marissa Reilly

